# Signaling interaction link prediction using deep graph neural networks integrating protein-protein interactions and omics data

**DOI:** 10.1101/2020.12.23.424230

**Authors:** Jiarui Feng, Amanda Zeng, Yixin Chen, Philip Payne, Fuhai Li

**Affiliations:** Institute for Informatics (I2), Washington University in St. Louis, St. Louis, MO, USA; Department of Pediatrics, Washington University School of Medicine, Washington University in St. Louis, St. Louis, MO, USA; Electrical and Systems Engineering Department, Washington University in St. Louis, St. Louis, MO, USA; Computer science, Washington University in St. Louis, St. Louis, MO, USA

## Abstract

Uncovering signaling links or cascades among proteins that potentially regulate tumor development and drug response is one of the most critical and challenging tasks in cancer molecular biology. Inhibition of the targets on the core signaling cascades can be effective as novel cancer treatment regimens. However, signaling cascades inference remains an open problem, and there is a lack of effective computational models. The widely used gene co-expression network (no-direct signaling cascades) and shortest-path based protein-protein interaction (PPI) network analysis (with too many interactions, and did not consider the sparsity of signaling cascades) were not specifically designed to predict the direct and sparse signaling cascades. To resolve the challenges, we proposed a novel deep learning model, *deepSignalingLinkNet*, to predict signaling cascades by integrating transcriptomics data and copy number data of a large set of cancer samples with the protein-protein interactions (PPIs) via a novel deep graph neural network model. Different from the existing models, the proposed deep learning model was trained using the curated KEGG signaling pathways to identify the informative omics and PPI topology features in the data-driven manner to predict the potential signaling cascades. The validation results indicated the feasibility of signaling cascade prediction using the proposed deep learning models. Moreover, the trained model can potentially predict the signaling cascades among the new proteins by transferring the learned patterns on the curated signaling pathways. The code was available at: https://github.com/fuhaililab/deepSignalingPathwayPrediction.

## 1. Introduction

A signaling pathway is defined as a set of molecular interactions within a cell that maintain or cause the change of specific biological processes of cells, like cell proliferation, migration, or differentiation. It is the critical components of the cellular machinery that convert the genotype and signals to specific phenotypes. One of the most critical objectives in cancer molecular biology is to uncover the causal genes and their signaling links or cascades among these candidate target genes that can potentially regulate tumor development and drug response [1]. A set of curated signaling pathways that play important roles in tumor cell development have been identified. For example, in the KEGG signaling pathway database, about 311 curated signaling pathways were collected. Moreover, the genetic mutations in differentially expressed genes in 10 cancer hallmark signaling pathways, like cell cycle, Hippo, Myc, Notch, NRF2, PI3K, RTK, RAS, MAPK, TGFbeta, P53, and beta-caenin[2], were investigated using the comprehensive multi-omics data of TCGA. The results indicated that 57% of TCGA cancer patients had at least one alteration on these signaling pathways. However, we still have a limited understanding of the complete picture of signaling pathways that regulate tumor cell development and their role in drug resistance. This is one major reason for the lack of effective treatments for cancer.

Signaling pathway or cascade inference is a rather challenging task due to the complex signaling pathways among large sets of genes within tumor cells. It remains an open problem. Many signaling network inference models are based on correlation, regression and Bayesian analysis [3]. For example, the weighted correlation network analysis (WGCNA) model was widely used[4]. However, it was designed to investigated the correlation, rather than the direct signaling cascade interactions. The correlation based models, like the BC3NET algorithm[5] and the SCINGE[6] models were proposed. The SCINGE model used kernel-based granger causality regressions predict the signaling cascades in pseudo temporally-ordered single-cell expression data. On the other hand, the protein-protein interaction (PPI) network was used in network analysis models based on the shortest paths of the up- or down-regulated genes. For example, the ARACNE model [7] was proposed to identify the master regulators on the activated signaling network based the transcription factors and target gene interactions.

Deep learning models have been reported to analyze the topological structures of PPIs mainly for the learning of node representations in the unsupervised learning. For example, DeepWalk[8] used a random walk to learn the latent representation of nodes and also applied social representation learning. The node2vec model [9] was proposed to embed the node into vector space. The SDNE[10] model used multi-layer non-linear functions to capture the network structure and generate node embedding. Also, the GNE model [11] used neural networks to encode gene expression, topological properties and expression information into dense vectors, which was similar to the Skip-gram[12] word embedding model. The experimental validation showed that GNE outperformed node2vec and LINE[13].

Recently, the deep graph neural network (GNN) [14] models have also been employed in the network analysis especially for the node representation analysis. For example, GraphSAGE[15] proposed an efficient way to learn the graph embedding inductively. GAT[16] applied the attention mechanism to actively learn a way to aggregate all the information in graphs. The DGCNN model [17] with sortPooling to efficiently learn graph features for graph classification. JKNet[18] adopted dense connection into GNN and used multi-hop massaging passing. Zhang et al[19] demonstrated that link information based on whole graphs can be approached by local graphs efficiently and proposed a GNN model called SEAL to conduct link prediction. Gu et al[20] extended graph attention networks in link prediction problems with fixed-sized neighborhood sampling to solve memory bottleneck and mini-batch problems during training.

However, the existing models were not been well designed to predict the sparse signaling cascades by integrating the omics data and topological structures of PPIs. For example, the directly interaction signaling cascades usually were ignored in the correlation, regression and Bayesian models because only a small set of genes have gene expression changes between normal and disease samples. Also, the shortest-path based protein-protein interaction (PPI) network analysis models cannot effectively identify the sparse signaling cascades from too candidate PPIs. To resolve these challenges, in this study, we proposed a novel deep GNN model, *deepSignalingLinkNet*, to predict signaling cascades between proteins. Specifically, the transcriptomics data and copy number data of a large set of cancer samples with the protein-protein interactions (PPIs) were integrated to identify the informative omics and PPI topology features in the data-driven manner to predict the potential signaling cascades by training the proposed model using the curated KEGG signaling pathways. Compared with PPI network, many genes have not been included in the curated KEGG signaling pathways. Thus, the trained model can potentially predict the signaling cascades among the new proteins by transferring the learned patterns on the curated signaling pathways.

## 2. Methodology

**Figure 1** shows the overview of the proposed model for signaling link prediction. The signaling link prediction problem can be formatted to predict if there is an interaction between two given genes or proteins. Specifically, the topological information were derived from the STRING[21] PPI database. Since the whole STRING PPI network has a large set of interactions among >17,000 genes or proteins, the one-hop neighbors of the given gene were identified to construct subgraphs: *G*(*V*_*a*_, *E*_*a*_) and *G*(*V*_*b*_, *E*_*b*_) for the genes *a* and *b*, where *V* and *E* represent the nodes and edges in the subgraphs. Then, the two subgraphs will be used in the PPIGE module to learn subgraph representations 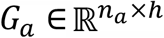 and 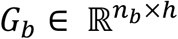, where *n*_*a*_ and *n*_*b*_ are the number of nodes in each graph and *h* is the hidden dimension in the model. The gene genomic data, i.e., gene expression and copy number variation, will be used in the GGE module to learn the genomic representations *h*_*a*_ ∈ ℝ^*h*^ and *h*_*b*_ ∈ ℝ^*h*^. Next, the GAGA module was proposed to learn a genomic aware gene topological representation, which aggregates the subgraph representation *G*_*a*_ and *G*_*b*_ based on genomic representations. The aggregation features are denoted as: *e*_*a*_ ∈ ℝ^*h*^ and *e*_*b*_ ∈ ℝ^*h*^ for the two genes respectively. Finally, *h*_*a*_, *h*_*b*_, *e*_*a*_, *e*_*b*_ are combined and used to compute the final prediction through the LED module. The detailed methodology was introduced in the following sections.

**Figure 1.**
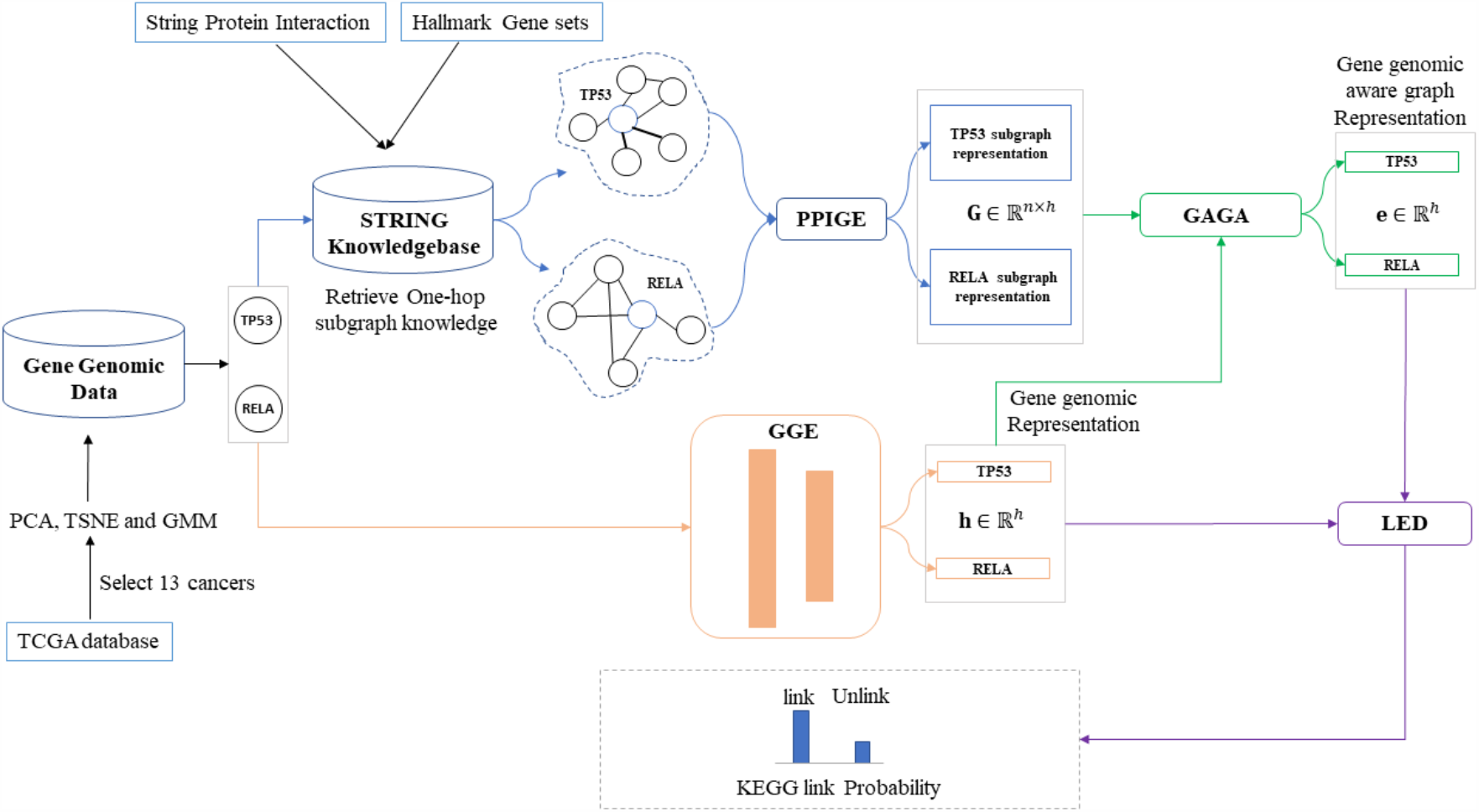
Overview of the deep signaling link prediction model.

### 2.1 Protein-Protein Interaction Graph Embedding Module (PPIGE)

To extract the topological information from the STRING PPI network, as proposed in [19], a *k*-hop subgraph was used to approximate the information of the whole graph efficiently. Specifically, we proposed the novel PPIGE module, which is a multi-layer graphical neural network with residual connections [22]–[24]. Let *X* ∈ ℝ^*n×m*^ be the initial representation of each node in the graph and *A* ∈ ℝ^*n×m*^ be the adjacency matrix of graph. Three versions of the PPIGE module were proposed based on the different type of GNN used: the GCN version, GAT version and self-attention version.

#### PPIGE-GCN version

The GCN version uses graph convolution networks [25] in each layer. If the input in each layer is *X*^*l*^, the representation of *X*^*l*+ 1^ is computed by:

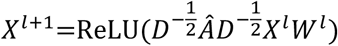

Where *Â* = (*A* + *I*), *I* is identity matrix. *D* is the degree matrix on *Â* and *W* is the learnable parameters matrix. In PPIGE-GCN, the number of GCN layers is 3. The first layer is *W* ∈ ℝ^*m×h*^ and the other two layers are *W* ∈ ℝ^*h*×*h*^. **Figure 2** illustrates the structure of the PPIGE-GCN version.

**Figure 2.**
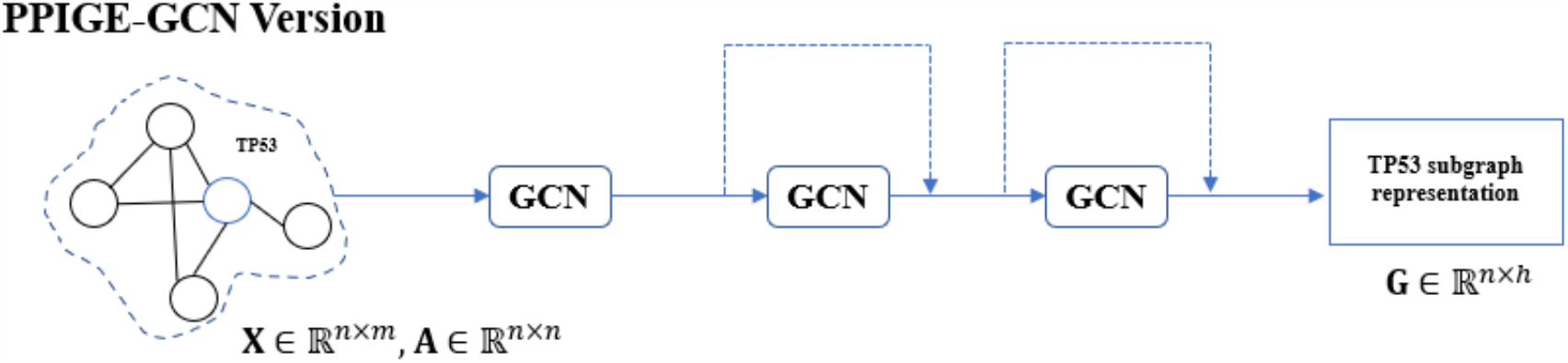
Structure of GCN version of PPIGE module.

#### PPIGE-GAT version

A limitation of GCN layers is that it sets the same weights of neighbors in convolution. To address this limitation, graph attention network(GAT)[16] was used to learn an attention weight for each node in graph. Let the input in each GAT layer be *X*^*l*^. The representation of *X*^*l*+ 1^ is computed by:

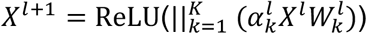

Where 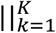 denotes the concatenation over *k* heads, *k* is the number of heads in multi-head attention, 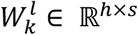 is the learnable parameters matrix. The attention score 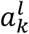 is computed by:

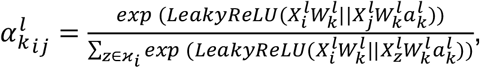

where 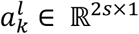 represents the learnable parameters and *ϰ*_*i*_ is the neighbors set for node 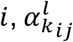 is the attention weight of neighbor node *j* to the node *i* in head *k* at layer *l*. To make the dimensions in each layer consistent, we set *s* = *h*/*k*. In PPIGE-GAT, there are two GAT layers. A linear projection layer is first used to convert the input dimension *m* to hidden size *h*. The architecture of the PPIGE-GAT version is shown in **Figure 3**.

**Figure 3.**
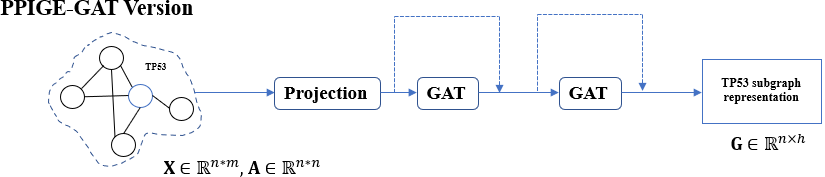
Structure of GAT version of PPIGE module

### 2.2 Gene Genomic Embedding Module (GGE)

For gene genomic data representation learning, the GGE module was introduced. GGE is a multi-layer neural network. For the gene genomic feature *x* ∈ ℝ^*ν*^, the computation in GGE is the following:

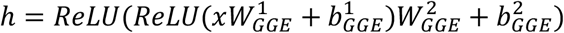

Where 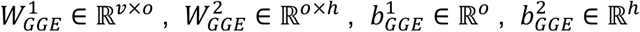 are learnable parameters matrixes. In the GGE model, we set *o* = 128, and *h* ∈ ℝ^*h*^ is the final genomic representation of the gene.

### 2.3 Gene Aware Graph Aggregation module (GAGA)

Problem specific aggregation functions, e.g., the mean aggregator or max aggregator, are important to learn the node representation. Herein, we proposed the GAGA module as follows. Let the *i-th* node in subgraph representation outputted by the PPIGEN module be *g*_i_ ∈ ℝ^*h*^, the gene genomic representation be *h* ∈ ℝ^*h*^. The GAGA aggregates the subgraph representation by:

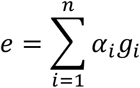

where *α*_*i*_ is the attention score for *i-th* node, which is computed by:

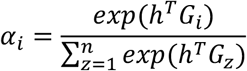

*α*_*i*_ characterizes how important a specific node in the subgraph for the node representation. The result *e* will be further used to predict the probability of the link between the source and target genes.

### 2.4 Link Existence Detector (LED)

With the genomic aware graph representation and the genomic representation, the probability of interaction link between two given genes is calculated using the LED module. First, the node representations are concatenated as follows:

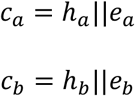

where || denotes concatenation operation, *c*_*a*_, *c*_*b*_ ∈ ℝ^2*h*^. Then the convolutional neural network (CNN) module was employed as:

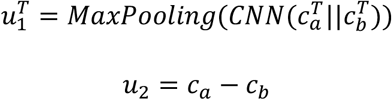

where 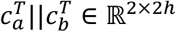. Then the signaling link prediction is calculated as:

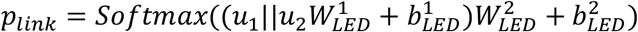

where the 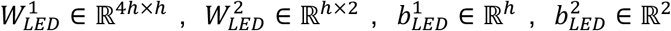 are the learnable parameters.

### 2.5 The Objective Function

To train the model, the focal loss[26] objective function was employed:

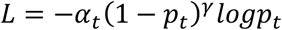

where *p*_*t*_ is the prediction of model for class *t. t* = 0 or 1 means that there is no signaling and there is a signaling link of two given genes respectively.

## 3. Results

In this section, we evaluated the performance of the deepSignalingLinkNet model, and showed that the proposed model outperformed other models in inferring KEGG signaling pathways.

### 3.1 Datasets

#### TCGA omics data and subtyping analysis

The Cancer Genome Atlas (TCGA) is a database that collected genomic data from 38 different types of cancer. The normalized gene expression data and copy number of TCGA samples were downloaded from the UCSC Xena data sever. Only the cancer types that had at least 10% normal samples or at least 20 normal samples were selected. In total, 13 different cancer types were kept.

The cancer samples were then clustered into 5 subgroups using the using the GMM algorithm on the dimensionally reduced gene expression dataset. The dimensions of the cancer samples to was reduced to 64 using the principle component analysis (PCA). Then, the dimension was further reduced to 2 using T-SNE[27] analysis. The subgrouping analysis was used to identify subtype-specific features, which might be missed by pooling all samples together. Then, the mean copy number and log2 fold change of the gene expression (between each subgroup and normal samples) of each group was constructedas the genomics features of each gene. This resulted in 130 features = 2(mean copy number and log2 fold change of gene expression) * 5 (subgroups of each cancer type) *13 (cancer types), for each of 18377 genes. The detailed number of samples in the selected TCGA cancer samples was listed in **Table 1**.

**Table 1.** \# of selected TCGA samples from 13 cancer types.

|  | # Control<br>sample | # Normal<br>sample | # G1 | # G2 | # G3 | # G4 | # G5 |
| --- | --- | --- | --- | --- | --- | --- | --- |
| **BRCA** | 1104 | 114 | 196 | 187 | 305 | 112 | 304 |
| **COAD** | 288 | 41 | 60 | 50 | 53 | 76 | 49 |
| **COADREAD** | 383 | 51 | 73 | 103 | 64 | 77 | 66 |
| **HNSC** | 522 | 44 | 127 | 116 | 100 | 106 | 73 |
| **KICH** | 66 | 25 | 14 | 9 | 4 | 23 | 16 |
| **KIRC** | 534 | 72 | 130 | 81 | 150 | 113 | 60 |
| **KIRP** | 291 | 32 | 71 | 44 | 90 | 20 | 66 |
| **LIHC** | 373 | 50 | 65 | 58 | 98 | 102 | 50 |
| **LUAD** | 517 | 59 | 105 | 125 | 69 | 144 | 74 |
| **LUNG** | 1019 | 110 | 220 | 72 | 204 | 273 | 250 |
| **PRAD** | 498 | 52 | 111 | 103 | 42 | 77 | 165 |
| **STAD** | 415 | 35 | 106 | 80 | 34 | 98 | 97 |
| **THCA** | 513 | 59 | 76 | 123 | 123 | 105 | 86 |

#### Hallmark Gene sets as features

[28]. Hallmark gene sets are a well curated gene sets that summarize specific, well-defined biological states or processes. In this study, the Hallmark gene sets were used as the initial gene features in the STRING database. The Hallmark Gene sets contain 50 different sets, which means that the dimension of gene feature is 50, and with 0 or 1 indicating if the gene belongs to that set. For genes that were not included in the Hallmark gene sets, their features were set as 0.

#### STRING PPI network

The STRING PPI network was collected from STRING database[29]. The 800 score threshold was used to filter out the low-confident PPIs, and then 17,179 genes and 841,027 signaling interaction links were obtained.

#### KEGG pathways

The training labels are obtained from the KEGG pathway database. KEGG (Kyoto Encyclopedia of Genes and Genomes) [30] is a database for the systematic understanding of gene functions. The KEGG signaling pathways provide knowledge of signaling transduction and cellular processes. There are a total 311 pathways. We collected links and the corresponding genes in all 311 pathways using the R package graphite. This resulted in a total of 8,002 genes and 116,820 links. Then, genes that did not have TCGA data or were not in STRING PPI network will be filtered out. Finally, 4347 genes and 70,431 signaling interaction links among 4,347 genes from 223 KEGG pathways. The details of all datasets are reported in **Table 2**.

**Table 2.** Statistics of Datasets

|  | # gene | # Link | # Feature |
| --- | --- | --- | --- |
| **STRING** | 17390 | 841027 | 0 |
| **KEGG pathway** | 8002 | 116820 | 0 |
| **Hallmark gene set** | 4383 | 0 | 50 |
| **TCGA** | 18377 | 0 | 130 |
| **Selected signaling interaction links** | 4347 | 70431 | 130 |

### 3.2 Model performance evaluation on KEGG signaling interaction links

In this evaluation test, 50% of the selected KEGG signaling interaction links were used as the training and testing positive data respectively. Then the same number of negative signaling interaction links (no interaction link) among the KEGG genes were selected randomly.

#### Model Setting

For the PPIGE models, the hidden output dimension *h* was set to 32. We also tested the GAT_large with *h = 64*. We added a dropout mechanism after each layer in the PPIGE and GGE module. The dropout rate was set to 0.1. The optimizer was Adam with *β*_1_ = 0.8, *β*_2_ = 0.999 and *ε* = 10^−7^. The learning rate for the three versions was 0.001. For each model, the weight decay was added with λ = 3 × 10^−7^. The epoch number was 30 for each model. The batch size was 64 for each model. Since the dataset is balanced, we set *α*_*t*_ = 1 and *γ*_*t*_ = 1 in focal loss, which make it equal to negative log likelihood loss. The gradient of the model was clipped with a threshold of 5.0 before each back-propagation step. The exponential moving average (EMA) was applied to all trainable variables with a decay rate of µ = 0.999. Let the model weights after the back-propagation step *t* be *W*_*t*_, and the model weights after the EMA in step *t* be *E*_*t*_. After back-propagation step *t* + 1, the new EMA model weights were updated with the function *E*_*t*+1_ = (1 − µ)*E*_*t*_ + *W*_*t*+1_. The model with highest accuracy in the test dataset was saved. The STRING subgraph knowledge is retrieved during training and testing. Due to computational and memory limitations, we randomly retrieved at most 200 neighboring nodes in the GCN modules. For GAT and GAT_large, at most 100 neighboring nodes were selected. The models were implemented by pytorch and trained on an MSI GeForce RTX 2070 GPU Super with 8Gb memory on a local machine. To compare the model performance using STRING PPI data alone and with the genomics data, the models without genomics data were named GCN_nonGeno and GAT_nonGeno. For each non-geno versions, the PPIGE module was the same as in the corresponding normal version, and only graph representation was used in the LED module to predict the distribution of link. Since the genomic data was removed, GAGA module was not be applied. The max aggregator [15] was applied to learn the final subgraph representation.

#### Model comparison

The multi-layer DNN, random forest, and kernel support vector machine (SVM) were selected for the comparison. For the multi-layer DNN, we used 4 layers and set the hidden sizes in each layer to be 128, 64, 32, 2 empirically. The activation function was ReLU for each layer except the final layer, which used the SoftMax function. The dropout mechanism was added after the first three layers. The dropout rate was 0.1. The optimizer was Adam with *β*_1_ = 0.8, *β*_2_ = 0.999 and *ε* = 10^−7^ and the learning rate was set to 0.001. For random forest and kernel-SVM, the input was the concatenation of the genomic data of two genes. For random forest, n_estimators and max_depth parameters were set to 80 and 7 respectively. For kernel-SVM, the radial basis function (RBF) kernel was used. The random forest and kernel-SVM were implemented using the scikit-learn package in python.

#### Evaluation results

Five evaluation metrics, accuracy, AUC, recall, precision and specificity, were used. **Table 3** showed the evaluation results. As seen, the proposed deepSignalingLinkNet outperformed other models. In addition, the integration of omics data with the PPI topological data improved the prediction accuracy. Moreover, the GAT (attention mechanism) version outperformed all other models. As seen in **Fig. 4**, the GAT had the similar performance compared with the GAT_large model, which were outperformed the GCN model significantly.

**Table 3.** Model comparison result

|  |  | Accuracy | AUC | Recall | Precision | Specificity |
| --- | --- | --- | --- | --- | --- | --- |
| Only genomic data | **RandomForest** | 0.714 | 0.798 | 0.721 | 0.714 | 0.708 |
|  | **Kernel-SVM** | 0.612 | 0.658 | 0.531 | 0.631 | 0.692 |
|  | **DNN** | 0.721 | 0.797 | 0.725 | 0.721 | 0.717 |
| Only topological data | **GCN_nonGeno** | 0.752 | 0.807 | 0.561 | **0.911** | **0.945** |
|  | **GAT_nonGeno** | 0.834 | 0.925 | 0.762 | 0.892 | 0.907 |
| The Proposed Model | **GCN** | 0.782 | 0.847 | 0.725 | 0.820 | 0.839 |
|  | **GAT** | **0.888** | **0.954** | **0.876** | 0.898 | 0.900 |
|  | **GAT_large** | 0.886 | 0.953 | 0.865 | 0.904 | 0.907 |

**Figure 4.**
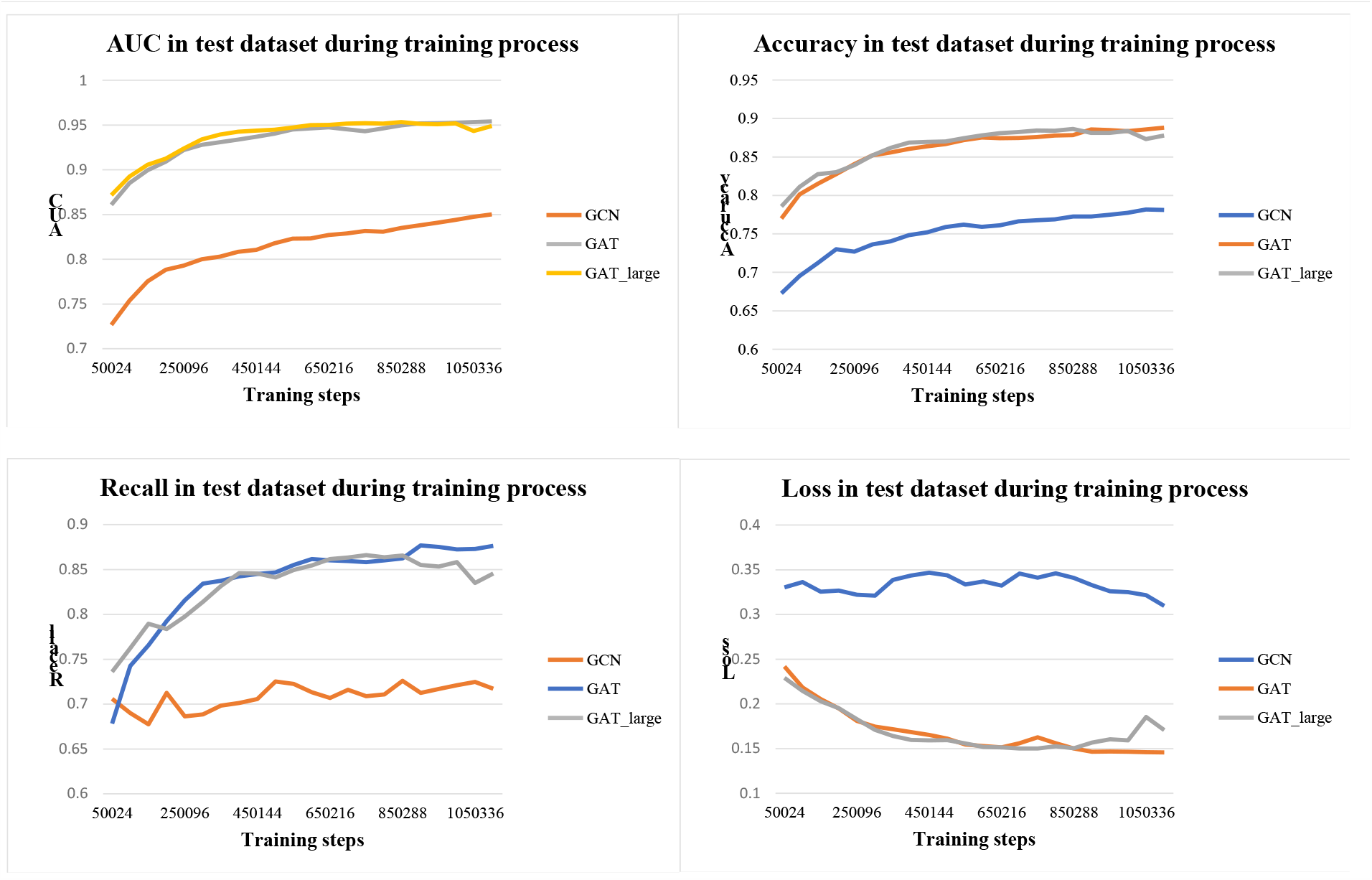
Trends of AUC, Accuracy, Recall and Loss of the GCN, GAT and AGT_large models.

### 3.3 Model performance evaluation on whole KEGG signaling pathways

In section 3.2, the models were evaluated by using randomly selected signaling interaction links from all the KEGG signaling pathways. In this section, the models were evaluated by using the whole signaling pathways, which can evaluate the transferability of the models to predict the signaling interaction links among a set of new genes or proteins.

#### Dataset Setting

All the signaling interactions of 10 pathways were selected as testing data. Then, 180 pathways were selected as training data. The remaining 33 pathways were used as validation data. For each signaling pathway in the testing and validation datasets, the signaling interactions were used as the true positive, and the rest gene pairs (without signaling interaction links) were used as true negative. For the training dataset, the signaling interactions were used as true positive, and 4 times number of true negative interactions were randomly selected.

#### Model Setting

Since had the best performance in the aforementioned validation, the GAT_large model was selected for this validation. All the configurations were the same as the above experiment except the epoch number was 10. We also set *γ*_*t*_ = 3 in focal loss considering the unbalanced training dataset. The four-fold cross-validation was employed to evaluate the models.

#### Evaluation results

**Table 4** showed the performance of the GAT_large version of the proposed model in test pathways. As seen, the average AUC of was ∼0.75, which was expected to be much lower than the AUC values in the above validation. One challenge is because that there were much more negatives than the true positive signaling interaction links. In another word, the curated signaling interaction links are much sparser compared with the STRING PPI interactions.

**Table 4.**
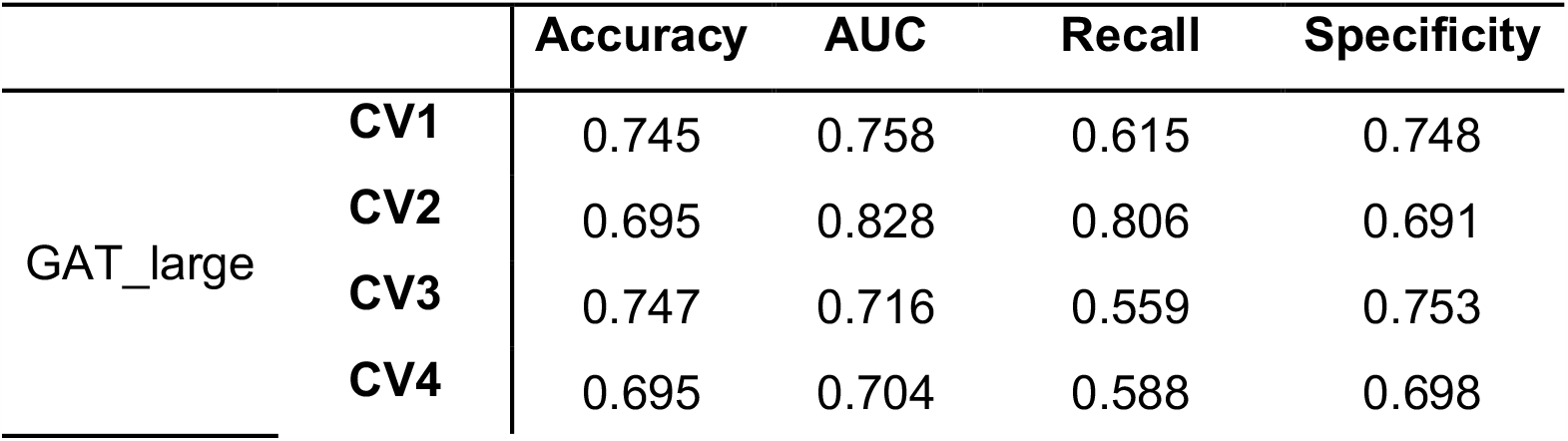
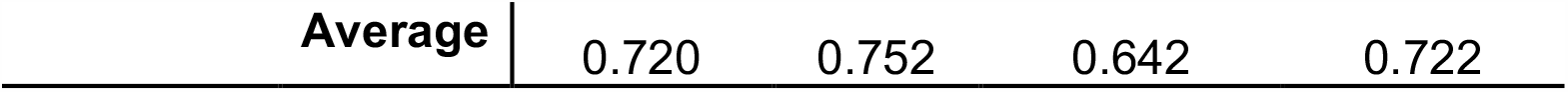
Signaling pathway inference result

## 4. Discussion and conclusion

Understanding the signaling cascades among a set of proteins that potentially regulate tumor development and drug response is one of the most critical and challenging tasks in cancer molecular biology. To date, there is a lack of effective computational models for the signaling interaction link prediction. Herein, we proposed a novel deep learning model, *deepSignalingLinkNet*, to predict signaling interaction links by integrating transcriptomics data and copy number data and PPIs. The advantages of the proposed deep learning model, compared with the gene co-expression and shortest path-based models, are that the model was trained using the curated KEGG signaling pathways to identify the informative omics and PPI topology features in the data-driven manner. The validation results indicated the feasibility of signaling cascade prediction using the proposed deep learning models. Also, it can potentially predict the novel signaling cascades among the new proteins by transferring the learned patterns on the curated signaling pathways.

This is a novel and exploratory study for signaling interaction link prediction. There are some limitations and challenges. For example, in the curated KEGG signaling pathways, there were only ∼4,000 proteins with the TCGA genomics data, whereas there were 17,000 genes in STRING PPI data. It is challenging to predict the new signaling interaction links of a large set new proteins that were not included in KEGG signaling pathways. Therefore, the integration of KEGG signaling pathways with other signaling pathway database, like wikiPathways[31], might be helpful to include more curate signaling pathways to train the model. Second, in addition to the Hallmark features of the genes, the Gene ontologies (GO) terms[32], and molecular gene set signatures[28] could also be helpful to annotate the functions of genes. Thirdly, the proposed model used the genomics data of 13 cancer types, which was developed for the general signaling interaction link prediction. It is interesting to expand the model to infer the patient-specific or disease subtype-specific activated signaling pathways like the gene co-expression and shortest-path based signaling network analysis models. We will investigate these challenges and limitations in the future work.

